# Proteomic Analysis of ACE Inhibitory Peptides extracted from Fermented Goat Milk

**DOI:** 10.1101/336107

**Authors:** Muhammad Zohaib Aslam, Sana Shoukat, Zhao Hongfei, Zhang Bolin

## Abstract

Protein extracted from goat milk was hydrolyzed with LH (Lactobacillus Helveticus-cicc22171). Angiotensin Converting Enzyme (ACE) inhibitory peptides were purified from fermented samples of goat milk protein with LH by optimizing incubation time to 8 hours (S-8), 16 hours (S-16), 24 hours (S-24) and 36 hours (S-36), via ultrafiltration. Molecular weight cut-off; 10000 Da (PM-10) membrane was used to perform size exclusion chromatography. Sample with 24 h incubation time was considered as best hydrolyzed as compared to others, by applying Nin-Hydrin reaction and SDS-PAGE analysis. ACE inhibitory assay validated the authenticity of S-24 in inhibiting ACE, in vitro. Furthermore, Q executive Hybrid Quadrapole-Orbitrap Mass Spectrometry was used to determine molecular structure and amino acid sequence of ACE inhibitory peptides. Two protein groups VLPVPQKAVPQ and VLPVPQKVVPQ containing PVP, VVP along with one most abundant peptide TQTPVVVPPFLQPEIMGVPKVKE containing VPP has been identified with highest ACE inhibitory activity on the basis of intensity, small structure and higher concentration of hydrophobic and aromatic amino acids. Fermented goat milk containing these novel bioactive peptides, can be used as nutraceuticals to inhibit ACE and control hypertension.

## 1. Introduction

Continual rise in blood pressure is defined as Hypertension (1). Hypertension is graded as the most repeatedly cardiovascular risk factor with the severity of 25–30% across the globe (2). Improper management of hypertension claimed seven million human lives along with 64-million life long disability worldwide (3). Genetic disposition, aging, overweight, lifestyle and nutrition can be considered as important factor of developing hypertension. Some diseases are also involved in creating hypertension, like Diabetes, Kidney disease, Pheochromocytoma, Cushing syndrome, congenital adrenal hyperplasia. Some medications like oral contraceptives, hormone therapy for menopause, and excessive intake of alcohol are contributor in developing hypertension along with other factors (4). Angiotensin I-converting enzyme (ACE; kininase II; EC 3.4.15.1) is considered as main root cause of hypertension that is carboxy-dipeptidyl-metallopeptidase and main linked enzyme of renin angiotensin system. It is involved in regulation of peripheral blood pressure. It mainly catalyzes the production of angiotensin-II from angiotensin-I (a vasoconstrictor) and inactivation of the vasodilator bradykinin (5, 6).

It has been known long ago about the importance of diet for human health. Recent studies concluded that diet is involved in the reduction of disease (7, 8). Milk protein is a bioactive component of milk. Along with energetic and nutritional value of milk protein, it serves as physiological regulator that is involved in reducing blood pressure by inhibiting the ACE (9-12). Studies confirmed that, individual with less milk consumption van have greater risk of hypertension. Many scientists have approved that milk proteins not only contain energetic and nutritional functions, but also can act as physiological regulators (9, 10). Scientists mainly focused on casein or whey protein to produce ACE inhibitory peptides; these peptides can be released within sequence of the parent protein and can be hydrolyzed by enzymes or food processing. Fermentation of milk also enhances its nutritional value through improved bioavailability of nutrients and production of substances that have a biological function (13). Antihypertensive peptides are the most, well studied among bioactive peptides. Bioactive peptides from bovine milk are not a topic of interest for most of the scientific community due to the extensive work. Now it is time to explore alternative sources like goat or sheep milk to extract bioactive peptides (14). Due to difference in casein contents, goat milk is considered better than cow milk (15).

This study is designed to evaluate the role of fermented goat milk (Guanzhong Goat) milk, hydrolyzed by LH in reducing hypertension with preconditions of optimizing incubation time along with identification and purification of ACE inhibitory peptides on the basis of peptide structure, intensity of peptides and Aromatic and hydrophobic peptide concentration.

## 2. Materials and Methods

### 2.1. Materials

Hippuryl-L-histidyl-L-leucine (Hip-His-Leu), purified rabbit lung ACE were purchased from Sigma Chemical Co. Ltd. Real Band 3-color high Range Protein Marker (9-245kDa) of reagent grade was purchased from BBI Life Sciences (Sangon Biotech Co. Ltd). Laboratory of Food Sciences (Beijing Forestry University) kindly donated stock culture of LH.

### 2.2. Ethics statement

All policies mentioned in the code of ethical committee (Beijing Forestry University) for the use and care of animals, were adopted and animal ethics committee of Beijing Forestry University approved the study design. Furthermore, Livestock department of Shaanxi province facilitated for milking process.

### 2.3 Milk Collection and Preparation of Samples

Fresh Goat milk from apparently healthy and uninfected guanzhong goats was collected from guanzhong area of Shanxi Province in China. Disinfectant solution (Safflon 20%) was used to wash the udder before milk collection. Autoclaveable plastic containers (1000 mL) were used for the collection of samples and sterilized at 121°C for 20 min (16). Whole milk was converted into skim milk by centrifugation at 4000 g for 10 min at 15°C and then stored at −20°C. Freeze-drying of skim goat milk was performed by Detianyou FD-1 Freeze Dryer (China Based Company). The freeze-dryer was programmed to operate at −50°C with a vacuum pressure of 20 pa (Pascal). After the end of freeze- drying cycle, dried milk was sealed under vacuum and stored at 5°C until further analysis (17).

Department of Food Sciences at Beijing Forestry University maintained stock culture of LH. Before experimental use, De Man, Rugosa and Sharpe (MRS) culture medium was prepared and stock culture was propagated for 16 to 18h at 37°C. Hemocytometer was used to calculate bacterial colonies and cells were harvested by centrifugation at 6000g for 10 min at 4 °C and washed three times with distilled water.

Goat milk Protein was co-cultured with the LH (1**×**10^9^cfu/mL) at the ratio of 10:2 mL respectively. Moreover, Samples were incubated for the interval of 8, 16, 24 and 36 hours. Furthermore, samples were centrifuged at 15000g for 30 minutes and supernatant was filtered by using 0.45µl (microliter) filter tip and stored at −20°C for further analysis.

### 2.4. Degree of hydrolysis

Nin-Hydrin reaction was applied to estimate the degree of hydrolysis of protein into peptides. To prepare 20mL solution Reagents were prepared according to standard protocol with 2g (gram) of Na_2_HPO_4_.10H_2_O, 1.2g NaH_2_PO_4_×2H_2_O, 0.1g Nin-Hydrin and 0.06g Fructose.

Each sample (S-8, S-16, S-24 and S-36) with 0.05mL quantity was added into 15ml of deionized water to dilute it further. Aliquot measuring in 0.5 mL was taken and mixed with 1ml reagent solution. Mixture was allowed to heat on 100°C for 15 minutes. After heating, the color of samples turned into purple and its wavelength is determined at 570nm by using Spectrophotometer (PGENERAL T-6 New century, China).

### 2.5. SDS-PAGE (sodium dodecyl sulfate–polyacrylamide gel electrophoresis)

Samples were dissolved into 50 mM Tris-HC1 with pH 6.8, containing 2% (w/v) SDS, 2 mM EDTA, 10% (v/v) glycerol and 0.1% (w/v) bromophenol blue. When dissociation was needed, 0.1% (w/v) dithiotreitot (DTT) was freshly added to the same buffer and the sample solution was heated at 95°C for 15 minutes. Polyacrylamide slab gels were cast and electrophoretic separations was carried out using a Bio-Rad Mini-Protein II Electrophoresis Cell System. Slab gel electrophoresis was performed in a 15% separating gel with a 5% stacking gel. Separating gel and stacking gels was prepared according to standard protocol.

Polyacrylamide gels were stained with 0.25% (w/v) Coomassie R-250 Blue in 50% (v/v) methanol and 7.5% (v/v) acetic acid for 15-30 min, according to discontinuous buffer system (18). Gel was destained in 20% (v/v) methanol and 7.5% acetic acid.

### 2.6. Size Exclusion Chromatography

Samples were filtered with 10000 Da cut-off membranes. Solution was further purified with gel filtration by using Sephadex G-15 column (1.6**×**90 cm) and eluted with deionized water. A Fraction of 5 mL from each vial was collected at the flow rate of 0.5 mL/min. Absorbance of fraction was calculated at 280nm by using spectrophotometer.

### 2.7. Assay for ACE inhibitory activity

ACE-inhibition activity of the goat milk peptides was measured according to (19) with some modification. Briefly, 180**μl** Of HHL buffer (5 mM Hip-His-Leu in 0.1 M borate buffer containing 0.3 M NaCl, pH 8.3) was mixed with 50**μl** of sample solution. The mixture of sample and buffer was incubated for 3 min at 37°C. Moreover, the reaction was started by the addition of 20μl of ACE (dissolved in distilled water, 0.1 units/ml). The whole mixture was incubated again for 40 min at 37°C. Adding 200μl of 1.0 N HCl has stopped the reaction. The absorbance was measured at 228 nm using a spectrophotometer to estimate ACE activity. The extent of inhibition was calculated as follows:

ACE inhibitor rate = (B-A/B-C) 100

Where A is the absorbance in the presence of ACE and with the ACE-inhibitory component, B is the absorbance with ACE and without the ACE-inhibitory component; C is the absorbance without ACE or ACE inhibitor component.

### 2.8. Identification of peptides

Q executive Hybrid Quadrapole-Orbitrap Mass Spectrometry was used to identify peptides in the sample. This is a bench top LC-MS/MS system combines quadruple precursor ion selection with high-resolution, accurate-mass Orbitrap detection; mass range from 50-6000 m/z and scan range up to 12 Hz. Chromatographic separations were performed on C18 column (300 μm i.d. × 5mm, packed with Acclaim PepMap RSLC C18, 5 μm, 100Å, Nano Viper, Acclaim PepMap 75 μm × 150mm, C18, 3 μm, 100A). The mobile phase consisted of 0.1% formic acid in pure water (Solvent A) and 0.1% formic acid, 80% ACN (Solvent B), formed with gradient elution with following parameters: 0-5 min, 0-5% B; 5-25 min, 5-5% B; 25-30 min, 5-50% B; 25-30%, 50-90% B; 30-35 min, 90% B; 35-45 min, 5% B. The total run time was 40 min with a constant flow rate of 300nL/min.

Some parameters in Orbitrap were as follows: spray voltage, 2.0 kV; capillary temperature, 250 °C; m/z (mass to charge ratio) range (ms), 350 to 1800. AGC (Automatic gain control) ion injection targets for first level FTMS (Fourier Transform Mass Spectrometry) scan were 70,000 (40 ms max injection time) with 3e6 AGS target and for second level mass spectrometry were 17,500 (60 ms max injection time) with 1e5 AGS target; 27NCE (normalized collisional energy) and 20 TopN.

### 2.9. Database Search

The original mass spectrum file was processed by MM File Conversion software. This PC (personal computer) version is a software suite for protein identification/characterization, cross-link/disulfide search, iTRAQ/TMT quantitation, and N/SILAC quantitation using LC-MS/MS to obtain the MGF format file, and then using MaxQuant searched the database. MaxQuant is one of the most frequently used platforms for mass-spectrometry (MS)-based proteomics data analysis.

## 3. Results and Discussion

### 3.1. Protein Degradation

In order to obtain Goat milk protein hydrolysates, LH were used to break down the protein into peptides. Degree of protein degradation into peptides by hydrolysis is called degree of hydrolysis. It is one of most authentic mechanism to measure the hydrolysis process (20). It is also used as indicator of comparing different protein hydrolysates (21). Protein concentration decreased with its degradation into peptides as the incubation time rises and maximum hydrolysis was observed after 24 hours. DH determined at various incubation times with LH is shown in Fig. 2.

**Figure 1.**
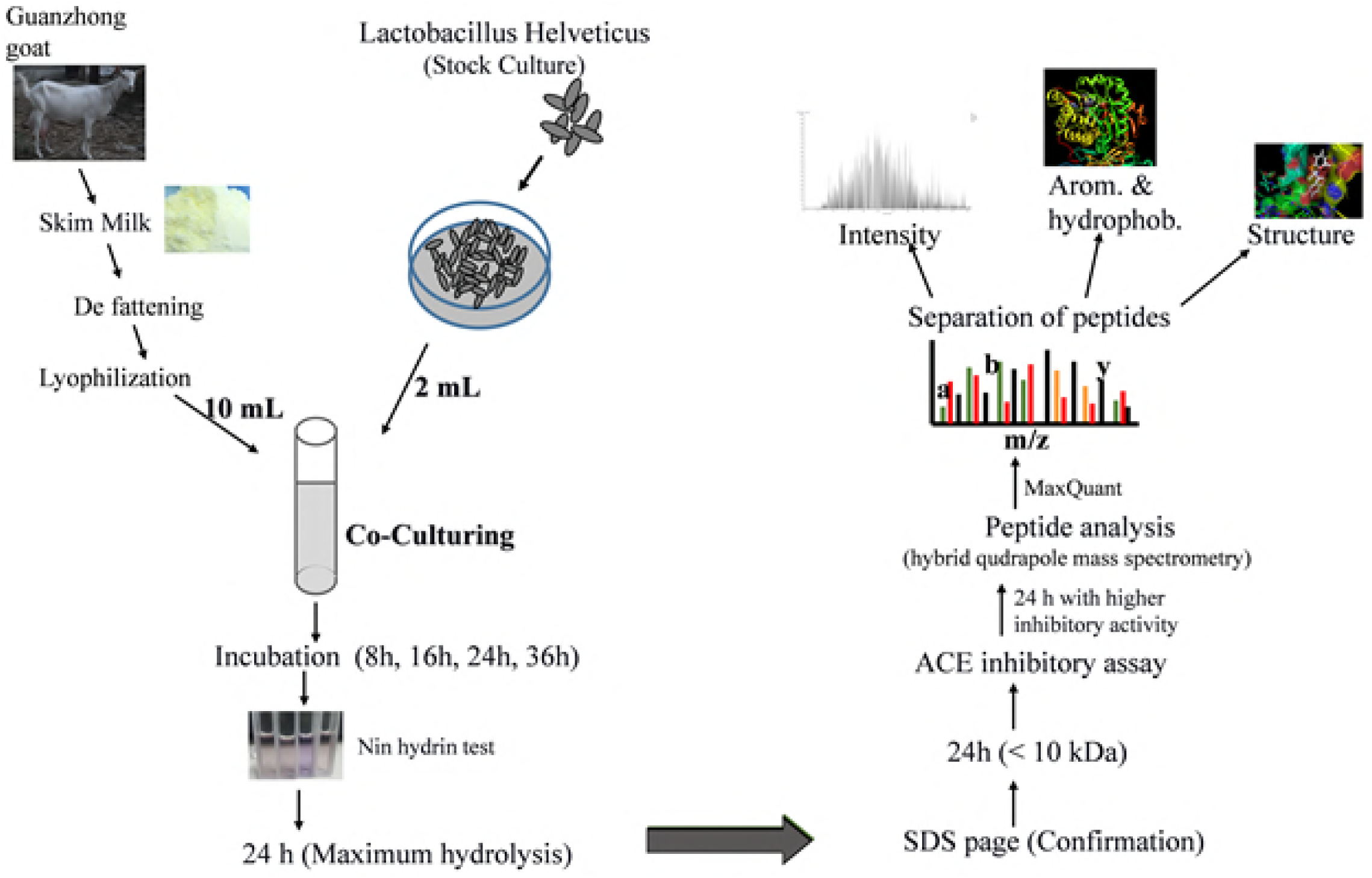
Graphical abstract of research.

**Figure 2.**
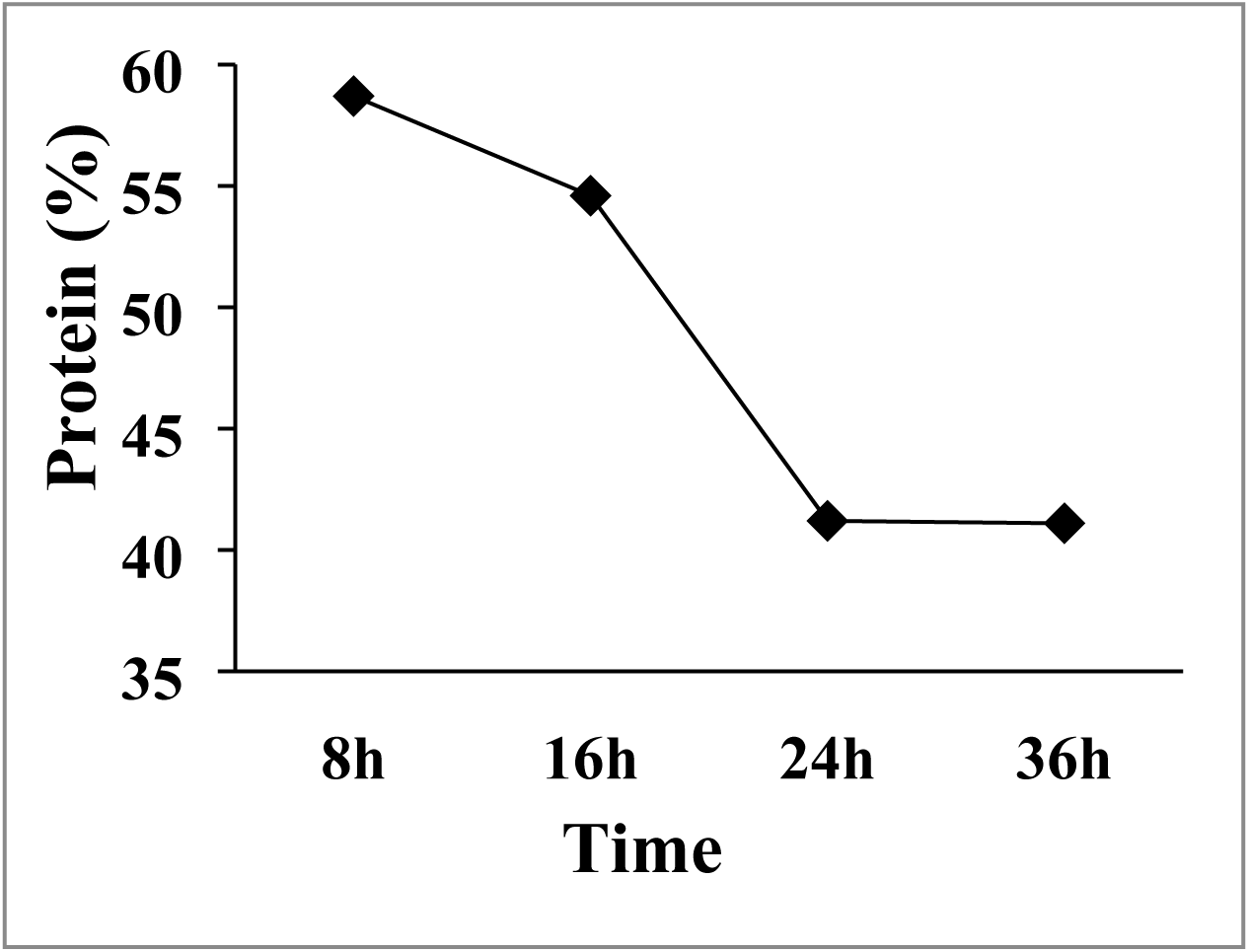
Time period was indicated in hours on x-axis while y-axis represents percentage of protein hydrolysis.

Results of DH show the gradual decrease in protein contents from 8 h to 24 and it remains almost constant from 24 to 36h, with negligible difference in protein degradation. It can be assumed on the basis of DH that sample should be fermented for 24 hours to get best hydrolysis rather than increasing time, because it can not make big difference in matter of peptides formation.

A representative SDS–PAGE electrophoregram of the samples with different incubation time is shown in Figure 3. This result endorses the outcome of Figure 2.

**Figure 3.**
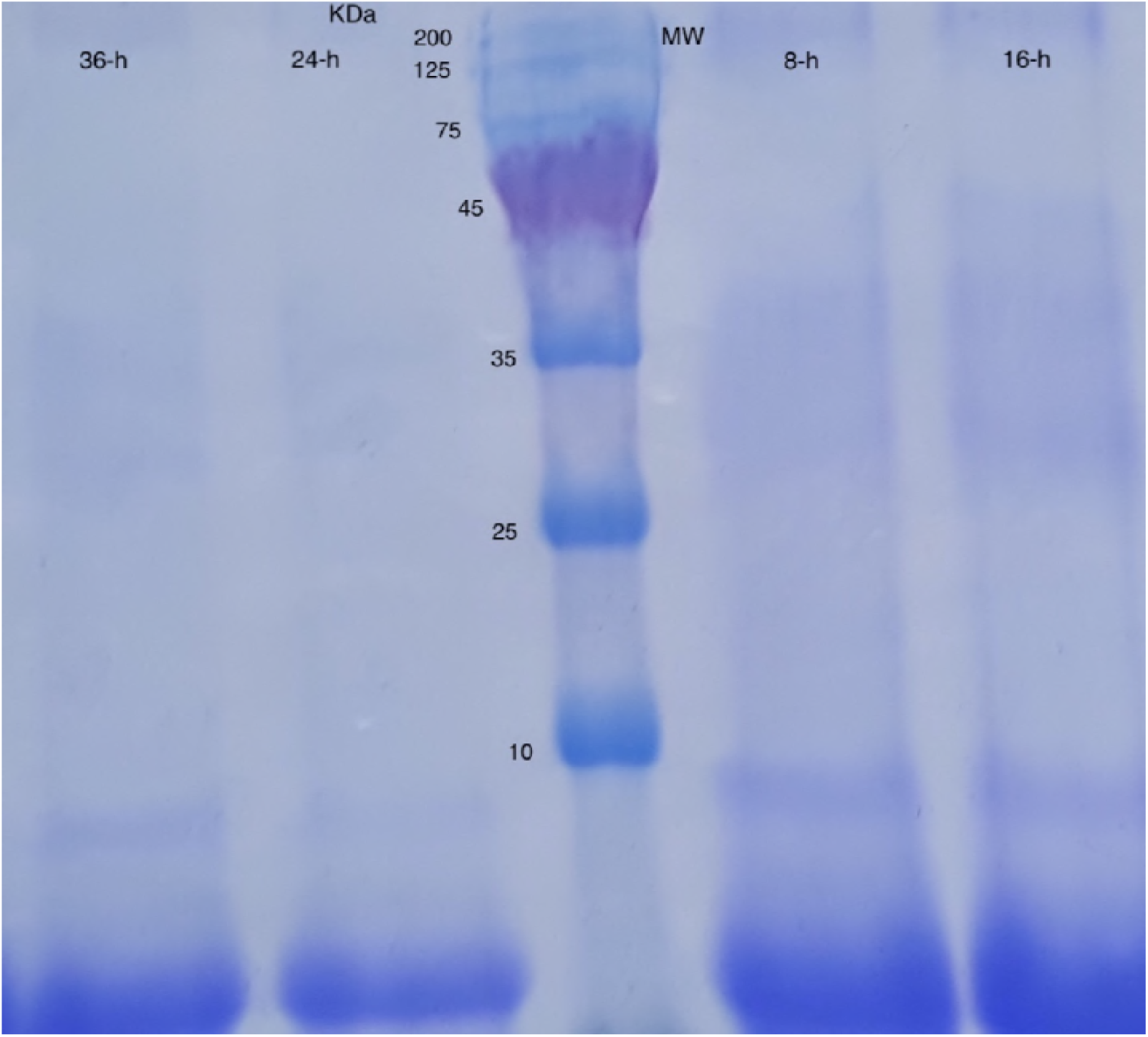
Molecular weight (MW) of peptides was indicated in kDa while samples with 8 h and 16 h incubation showed the presence of 10kDa peptides while samples with 24 h and 36-h incubation period showed absence of 10kDa peptides.

Major changes while performing SDS-PAGE included decrease in several bands indicated greater enzymatic breakdown by 24-h incubation. Furthermore, 10% acrylamide gel was used to identify the Small molecular parts. Samples with 24 h and 36 h incubation time, represented almost same picture of hydrolysis that indicated importance of 24-h incubation sample.

### 3.2. Ultrafiltration

Filtration of samples with PM-10 membrane (molecular weight cut-off; 10 kDa) was done to further purify ACE inhibitory peptides. Samples were lyophilized after purification and concentrated in distilled water.

Different steps in separation and purification techniques were recommended (22). For this purpose, Sephadex G-25 column was used to load the sample and peptides with low molecular mass (under 3000 Da) were separated. It indicates the role of small peptides in obstruction of ACE and these results were in agreement with the previous work (23, 24).

### 3.3. ACE Inhibition Pattern

All four samples were tested to evaluate their potency to inhibit ACE production by applying ACE assay, in vitro. Although, It was confirmed that S-24 was best hydrolyzed by previous tests and expected to show better inhibitory activity due to greater concentration of hydrolyzed protein, but all samples were tested against ACE to further authenticate results. A clear trend of increasing ACE inhibitory activity has been observed with increasing incubation time of fermented samples, and after 24 hours, increasing trend was as slow as negligible. It was clear after running assay that S-24 was the sample to further evaluate for its peptides profile that were showing greater ACE inhibitory activity as compared to others. Figure 4, indicates the increasing trend of ACE inhibition percentage with increasing time. After 8 h of incubation 51.11% ACE inhibitory percentage was observed that increased gradually with time and after 16-h, it rose to 62.23%. Further increase in time to incubate samples, resulted in increase in inhibition of ACE by 72.32% after 24h and 73.98% after 36h.

**Figure 4.**
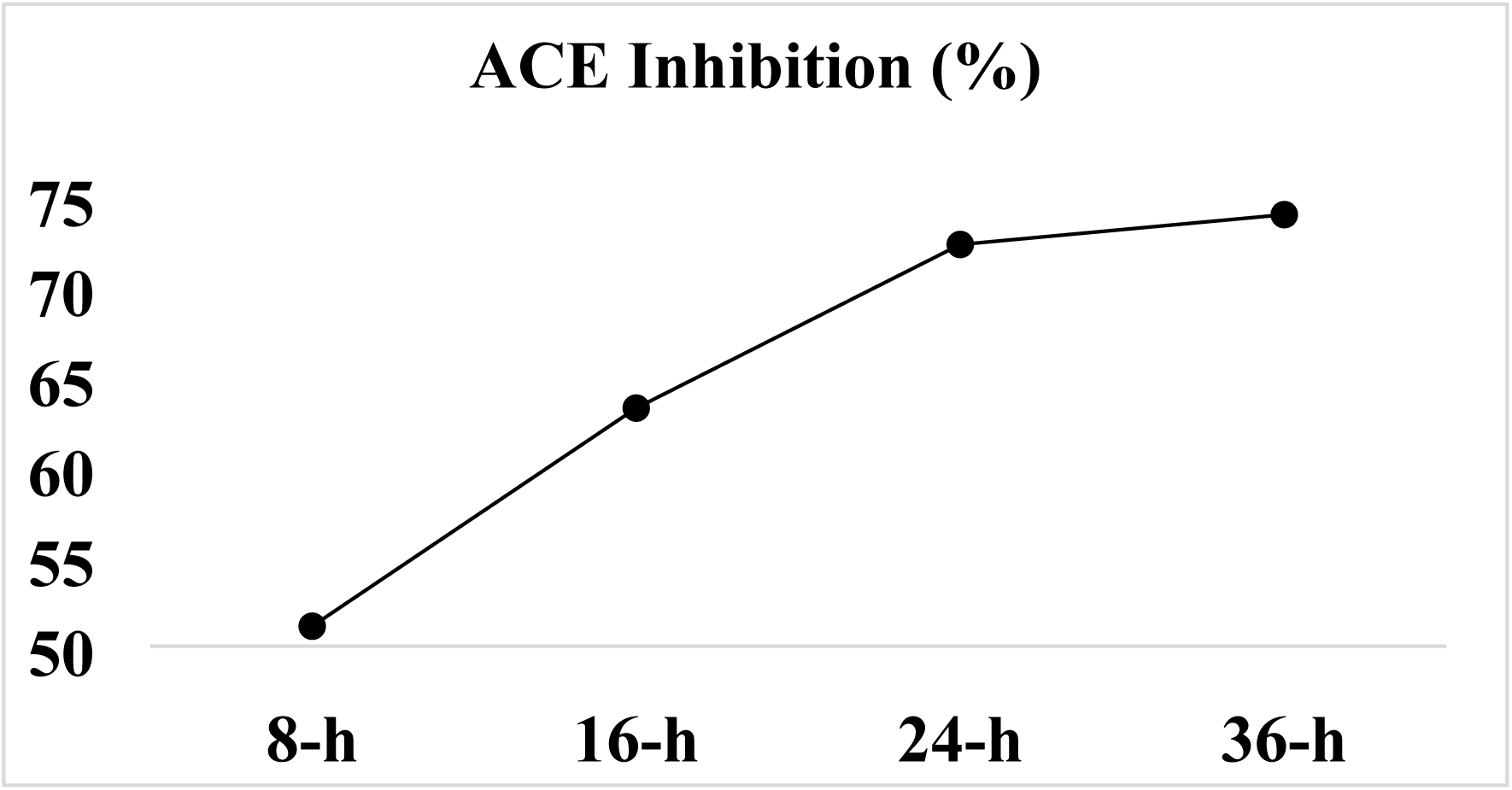
Time period in hours, is indicated on x-axis while Y-axis represents the level of ACE inhibitory percentage.

### 3.4. Identification of Peptides

Five most abundant Peptides with two unique protein groups have been identified as shown in table 1.

**Table 1.** Peptide ID was assigned on the basis of MaxQuant search results, and Peptide sequence was obtained in the result of running MS/MS.

| Peptide ID | Peptide Sequences | Mass (Da) | Intensity (10<sup>8</sup>) | C-term cleavage window | N-term cleavage window |
| --- | --- | --- | --- | --- | --- |
| 1073 | TQTPVV<u>V</u>PPFLQPEIM<br>GVPKVKE | 2548.397 | 3.46 | MGVPKVKETM<br>VPKHKE | LPQNILPLT<br>QTPVVVP |
| 1045 | TLTDVEKL | 917.506 | 3.13 | TLTDVEKLHLPL<br>PLVQ | PFTESQSLT<br>LTDVEKL |
| 1136 | VLP<u>V</u>PQKAVPQ | 1174.707 | 3.01 | VPQKAVPQRDM<br>PIQAF | VLSLSQPK<br>VLPVPQKA |
| 826 | REQEELNVVGE | 1300.625 | 2.98 | EELNVVGETVE<br>SLSSS | CLVALAIA<br>REQEELNV |
| 1138 | VLP<u>V</u>PQK<u>V</u>VPQ | 1202.738 | 2.72 | VPQKVVPQRDM<br>PIQAF | VLSLSQPK<br>VLPVPQKV |

They were categorized on the basis of intensity and abundance of peptide in the sample, presence of hydrophobic and aromatic amino acids and structure of amino acid sequence. Peptide sequence, with molecular weight of 2532.4026 Da and PID-1073 (peptide identification number), was categorized as the most abundant peptide on the basis of Mass to Charge ratio (m/z) which was recorded 1267.72 in figure 5 (A). It was identified several times with retention time of 41.785, 41.824, 44.933 and 44.944 min. Low molecular weight peptide (917.506 Da) with the retention time of 25.411, 27.137 and 27.93 min was identified thrice in S-24. Mass spectrum of PID-1045 was 459.76 m/z shown in figure 5 (B). Third and fifth peptides with PVP, VVP sequence contested for potential ACE inhibitor peptides. PID-1136 with two times spectroscopic identification and retention time of 21.494 and 21.519 min while PID-1138 with one time identification with retention time of 25.318 min, was explained in figure 5 (C) and 5 (E). Molecular weight of PID-1136 and 1138 was 1174.707 Da and 1202.738 Da respectively. Within Unique protein groups, PID-1136 with leading razor protein number Q712N8 was identified after 35 scans containing 6 isotopic peaks while PID 1138 with leading razor protein number Q95L76, was identified after 52 scans containing 5 isotopic peaks. Furthermore, PID 826 was identified once with retention time of 21.648 min. Its molecular weight was 1300.625 Da explained spectroscopically in figure 5 (D). PID 826 with leading razor protein number Q95L76 was identified once after 26 scans containing 6 isotopic peaks.

**Figure 5(A), (B), (C), (D) and (E).**
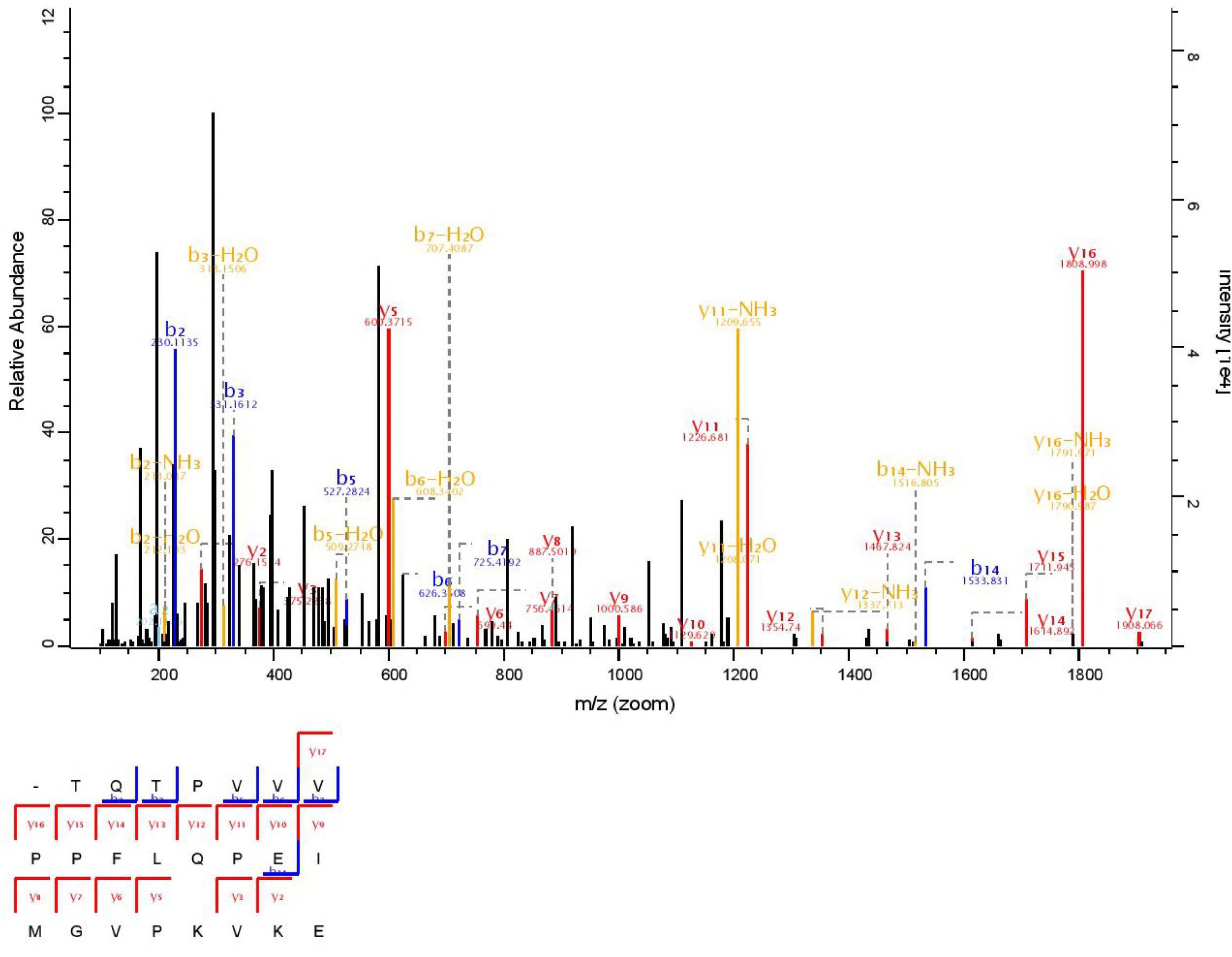
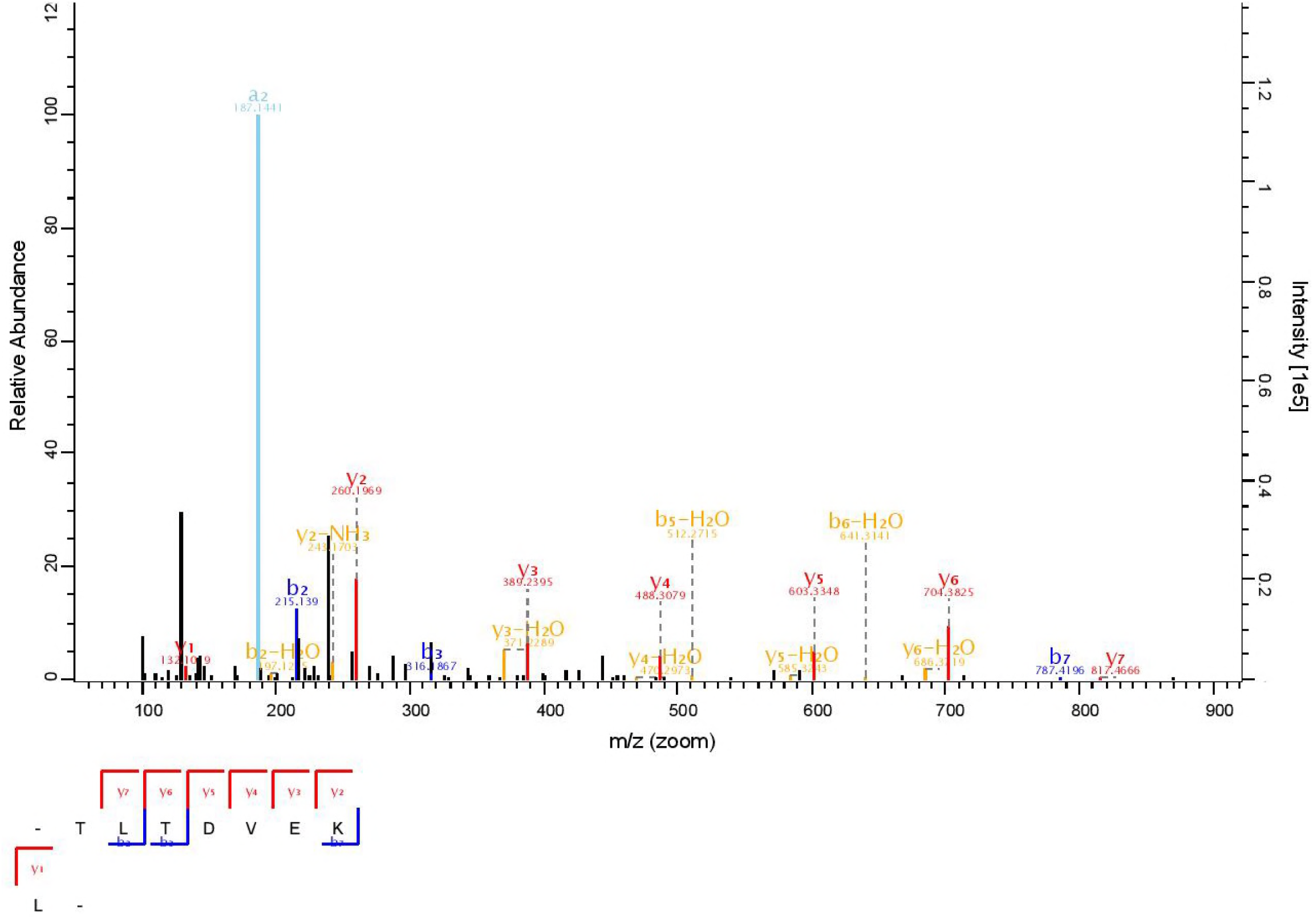
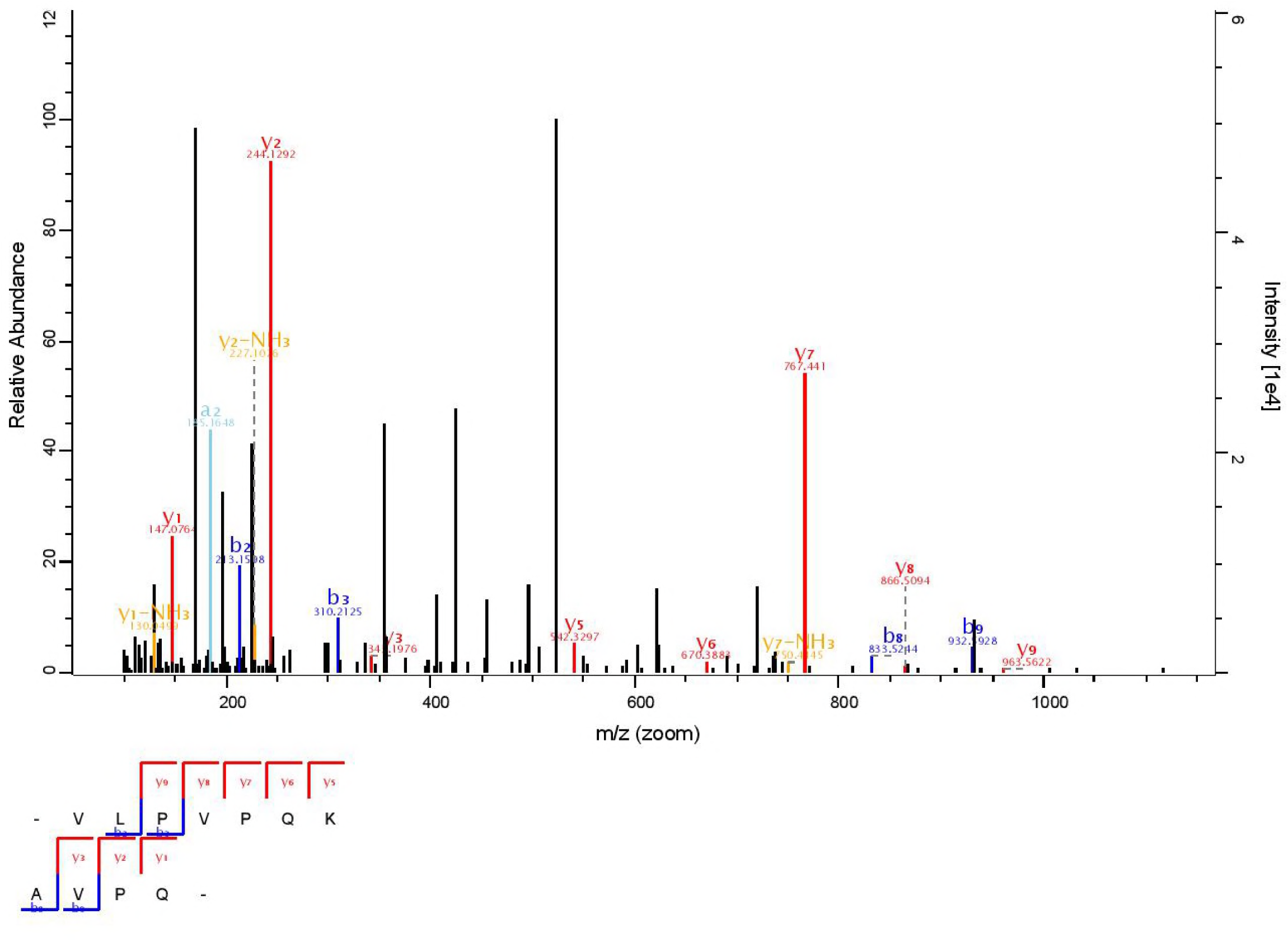
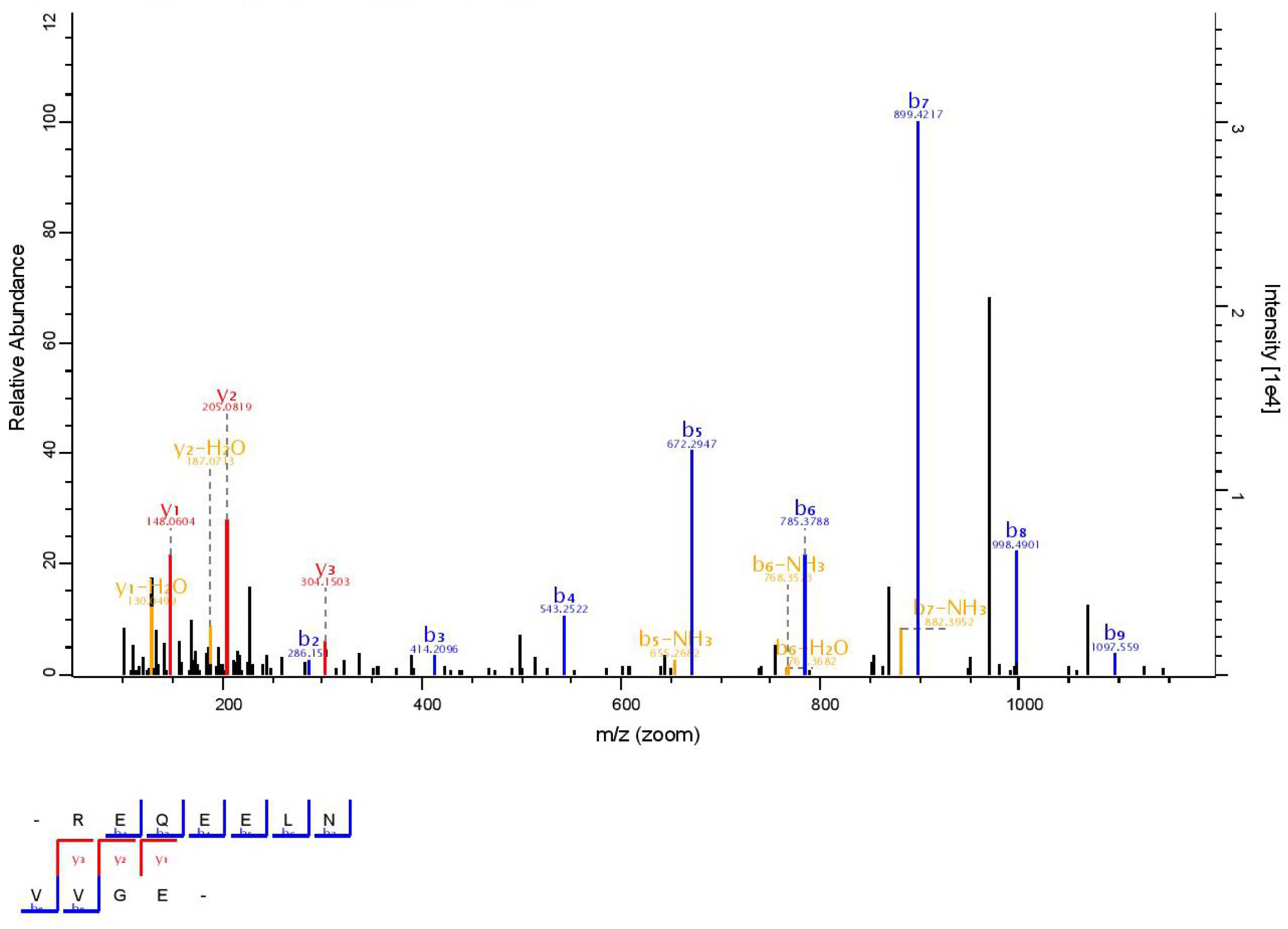
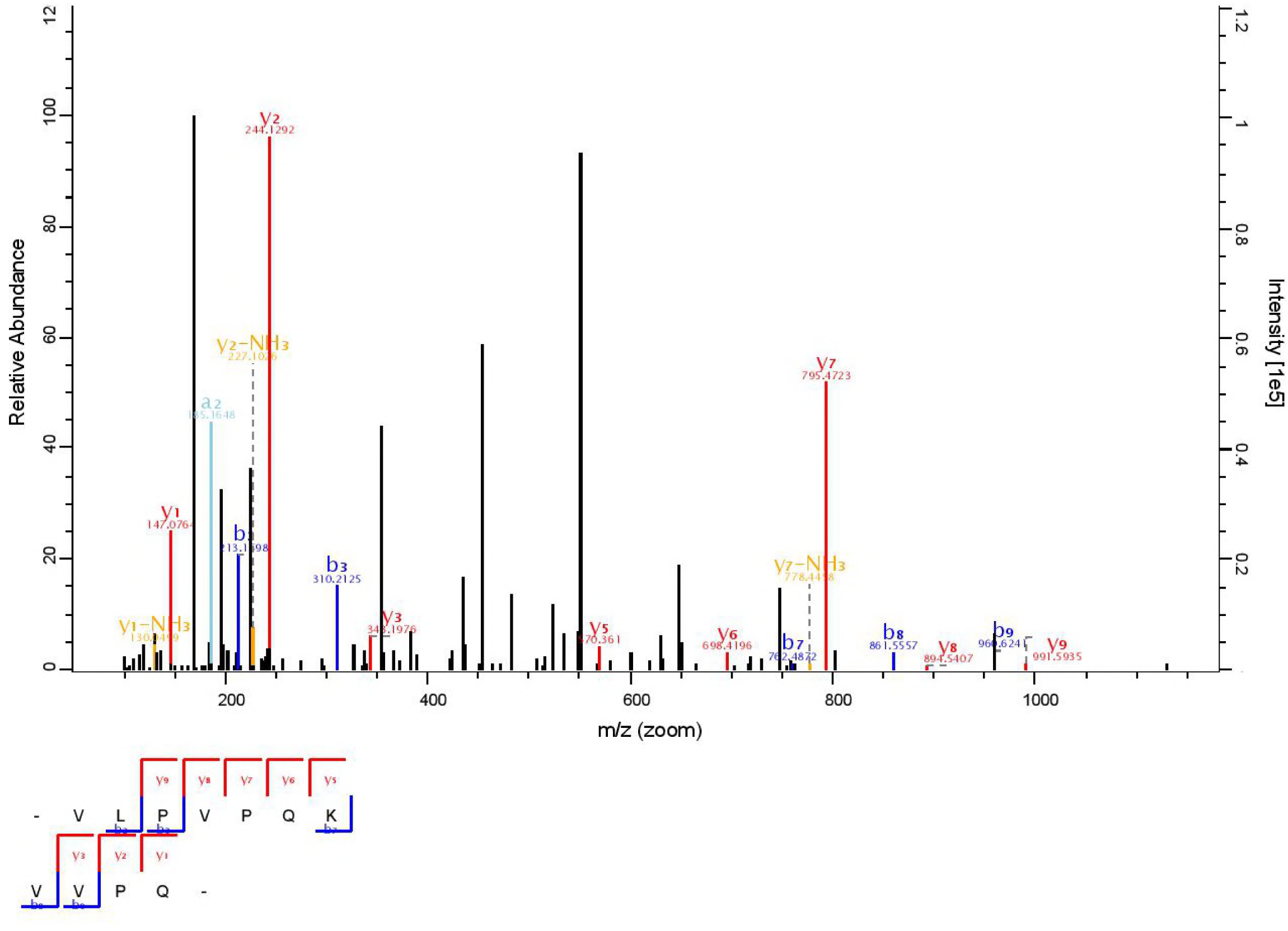
The MS/MS spectrum of charged ions with m/z 1267.72 (TQTPVVVPPFLQPEIMGVPKVKE), 459.76 (TLTDVEKL), 588.36 (VLPVPQKAVPQ), 651.32 (REQEELNVVGE) and 602.38 (VLPVPQKVVPQ) have been identified (Figure 5 (A), 5 (B), 5 (C), 5 (D) and 5 (E). The MS/MS spectrum was acquired with Q executive Hybrid Quadrapole-Orbitrap Mass Spectrometery. The peptide sequence is indicated on the bottom with collision-induced fragmentation pattern. The b and y ions are indicated in blue and red, respectively. X-axis shows the m/z and y-axis shows the relative abundance of each amino acid.

It was reported earlier that many ACE inhibitory peptides consist of 2–12 amino acid residues (25), and our samples indicates that out of five most abundant peptides, Four (TLTDVEKL, VLPVPQKAVPQ, REQEELNVVGE, VLPVPQKVVPQ) consist of less than 12 peptides with low molecular weight, which is in accordance to our findings. Short chain peptides are less prone to proteolytic enzymes of small intestine after oral administration, while easily absorbable to the absorptive cells because intestinal cells are restricted to absorb ultra-small peptides (26). While an other study was conducted by running experiments on rats and concluded that peptides larger than ultra-small peptides can also passed through small intestine wall but absorption mechanism of larger peptides is not yet explained properly that needs to be explored (27). On the other hand, peptides detected in S-24 showed ACE inhibitory activity, in vitro and it might assume that these peptides will remain intact in the gut and intestine of human because of their smaller structure with low molecular weight rather than degrading by proteolytic enzymes and might be absorbed easily through intestinal cells. It was reported in 2004 that digestive enzymes are unable to break down some peptides like Proline- and hydroxyproline-containing peptides and tripeptides containing the C-terminal proline-proline due to resistance of those peptides (28), while higher concentration of Proline in PID 1073, 1136 and 1138 respectively was identified in this study that can by-pass digestive enzymes easily and make them active contestant in inhibiting ACE. Besides, peptide size, there are other factors like amino acid residues and peptide sequence, may play a significant role in shaping ACE-inhibitory activities. It is stated in some studies that ACE-inhibitory peptides encoded in natural foods and products contain Proline, Lysine or aromatic amino acid residues (29, 30) that further supports our finding by exhibiting the presence of Lysine in all peptides with higher presence of Proline in PID 1073 followed by PID 1136 and 1138. Greater quantity of hydrophobic aliphatic amino acid and aromatic amino acid like Proline, Leucine, Alanine, Metionine and Isoleucine, in a sample may increase the ACE-inhibitory activity (31-33), While our findings are in agreement by the identification of peptides like TQTPVVVPPFLQPEIMGVPKVKE and TLTDVEKL with higher concentration of aromatic and hydrophobic amino acid. Methiotine, Triptophane, and Valine in P-1073, 1045, 1136 and 1138 on C-cleavage indicates the greater hydrophobicity of this peptide and greater hydrophobicity of C-cleavage shows the greater ACE inhibitory activity (34). C-cleavage with Aromatic amino acid and N- cleavage with Aliphatic amino acid might increase ACE inhibitory activity (35), While in this study, an aromatic amino acid Tyrocine was identified in P-1045 and aliphatic amino acid Leucine, Valine and Valine was identified in PID-1073, 1136 and 1138 respectively. Leucine and Valine, with small N-terminal amino acids and strong hydrophobicity are more appropriate for ACE inhibitory activity (36) and same amino acids were found on N-terminal of P-1073, 1136 and 1138.

## 4. Conclusion

The molecular mass of peptides (peptide length) is not the only important factor to evaluate its functional activities, other factors including presence and intensity of aromatic and hydrophobic amino acid and amino acid sequence also has an important role to play. In this study, fermented goat milk peptides, by optimizing incubation time to 24 hours, were proved to exhibit remarkable ACE-inhibitory activity. Three ACE-inhibitory peptides, PID-1073 (TQTPVVVPPFLQPEIMGVPKVKE), PID-1136 (VLPVPQKAVPQ) and 1138 (VLPVPQKVVPQ) were identified by Q executive Hybrid Quadrapole-Orbitrap Mass Spectrometery from S-24. Furthermore, structure analysis, intensity and presence of hydrophobic/aromatic amino acid concentration provided clear picture of their mode of action. It provided a basis to synthesize these peptides artificially. Goat milk peptides fermented with LH after optimizing incubation time to 24 hours can be used as functional food or nutraceuticals in treating hypertension.

## Acknowledgement

I am really thankful to the Zhao Hongfei, Lecturer (Beijing Forestry University, for providing me all opportunities to run experiments. This research was done with funding of Beijing Key Laboratory of Forest Food Processing and Safety, School of Biological Sciences and Technology, Beijing Forestry University (China).

## Conflict of Interest

I declare no conflict of interest.

